# Neurobehavioural Correlates of Obesity are Largely Heritable

**DOI:** 10.1101/204917

**Authors:** Uku Vainik, Travis Baker, Mahsa Dadar, Yashar Zeighami, Andréanne Michaud, Yu Zhang, José C. García Alanis, Bratislav Misic, D. Louis Collins, Alain Dagher

## Abstract

Recent molecular genetic studies have shown that the majority of genes associated with obesity are expressed in the central nervous system. Obesity has also been associated with neurobehavioural factors such as brain morphology, cognitive performance, and personality. Here, we tested whether these neurobehavioural factors were associated with the heritable variance in obesity measured by body mass index (BMI) in the Human Connectome Project (N=895 siblings). Phenotypically, cortical thickness findings supported the “right brain hypothesis” for obesity. Namely, increased BMI associated with decreased cortical thickness in right frontal lobe and increased thickness in the left frontal lobe, notably in lateral prefrontal cortex. In addition, lower thickness and volume in entorhinal-parahippocampal structures, and increased thickness in parietal-occipital structures in participants with higher BMI supported the role of visuospatial function in obesity. Brain morphometry results were supported by cognitive tests, which outlined a negative association between BMI and visuospatial function, verbal episodic memory, impulsivity, and cognitive flexibility. Personality-BMI correlations were inconsistent. We then aggregated the effects for each neurobehavioural factor for a behavioural genetics analysis and estimated each factor’s genetic overlap with BMI. Cognitive test scores and brain morphometry had 0.25 - 0.45 genetic correlations with BMI, and the phenotypic correlations with BMI were 77-89% explained by genetic factors. Neurobehavioural factors also had some genetic overlap with each other. In summary, obesity as measured by BMI has considerable genetic overlap with brain and cognitive measures. This supports the theory that obesity is inherited via brain function, and may inform intervention strategies.

**Significance Statement:** Obesity is a widespread heritable health condition. Evidence from psychology, cognitive neuroscience, and genetics has proposed links between obesity and the brain. The current study tested whether the heritable variance in body mass index (BMI) is explained by brain and behavioural factors in a large brain imaging cohort that included multiple related individuals. We found that the heritable variance in BMI had genetic correlations 0.25 - 0.45 with cognitive tests, cortical thickness, and regional brain volume. In particular, BMI was associated with frontal lobe asymmetry and differences in temporal-parietal perceptual systems. Further, we found genetic overlap between certain brain and behavioural factors. In summary, the genetic vulnerability to BMI is expressed in the brain. This may inform intervention strategies.

## Introduction

Obesity is a widespread condition leading to increased mortality (1) and economic costs (2). Twin and family studies have shown that individual differences in obesity are largely explained by genetic variance (3). Gene enrichment patterns suggest that obesity-related genes are preferentially expressed in the brain (4). While it is unclear how these brain-expressed genes lead to obesity, several lines of research show that neural, cognitive, and personality differences have a role in vulnerability to obesity (5, 6). Here we seek to test whether these neurobehavioural factors could explain the genetic variance in obesity.

In the personality literature, obesity is most often negatively associated with Conscientiousness (self-discipline and orderliness) and positively with Neuroticism (a tendency towards negative affect) (7). In the cognitive domain, tests capturing executive function, inhibition, and attentional control have a negative association with obesity (5–8). Neuroanatomically, obesity seems to have a negative association with the grey matter volume of prefrontal cortex, and to a lesser extent the volume of parietal and temporal lobes, as measured by voxel based morphometry (9). It has also been suggested that structural and functional asymmetry of the prefrontal cortex might underlie overeating and obesity (10). For genetic analysis, cortical thickness estimates of brain structure from Magnetic Resonance Imaging (MRI) have been preferred over volumetric measures (11). However, to date, reports of cortical thickness patterns associated with obesity have been inconsistent (12, 13). As a prerequisite to our goal of ascertaining the heritability of brain-based vulnerability to obesity, we sought to extend previous neurobehavioural findings in a large multi-factor dataset from the Human Connectome Project (HCP). We also measured volumetric estimates of medial temporal lobe and subcortical structures, which have been implicated in appetitive control (e.g., 14).

The main goal was to assess whether the aforementioned obesity-neurobehavioural associations are of genetic or environmental origin. Recent evidence from behavioural and molecular genetics suggests that there is considerable genetic overlap between obesity, cognitive test scores, and brain imaging findings (15–20). However, the evidence so far is not comprehensive across all neurobehavioural factors discussed. A recent paper assessed the heritability of obesity-associated regional brain volumes (21). However, the study did not analyze the heritability of the association between brain and obesity. The latter analysis is crucial for understanding whether brain anatomy and obesity could have a genetic overlap, which would suggest that the heritability of vulnerability to obesity is expressed in the brain.

In addition, we sought to estimate the genetic overlap between the different BMI-related neurobehavioural factors. On one hand, performance on cognitive tests and personality must originate from the brain (e.g., 22), and therefore personality and cognition could be expected to explain brain-morphometry associations with BMI (6). On the other hand, brain-behaviour associations are far from certain (23), and even different measurement traditions in both behaviour (personality and cognitive tests) and brain morphometry (cortical thickness or brain volume) are often conceptualized as providing independent sources of information (7, 11). Documenting the degree of genetic overlap between behavioural and brain measures would shed light on whether similar underlying processes lead to obesity’s associations with different neurobehavioural factors.

Taken together, the goal of the current analysis was to use a large multifactor dataset to analyze the heritability of the associations between obesity and brain/behaviour. We further tested genetic overlap between the different neurobehavioural factors themselves.

## Results

### Background

We analyzed data from 895 participants from the Human Connectome Project S900 release (24), including 111 pairs of monozygotic twins and 188 pairs of dizygotic twins and siblings. Similarly to many previous reports (3) we modelled BMI heritability with the AE model (A: additive genetics and E: unique environment), as opposed to the ACE model (C: common environment), as AE had the lowest Akaike Information Criterion (Dataset S1, section 9). BMI heritability was A=71% [95% CI: 61%;78%], which is close to the published meta-analytic estimate (A=75%, 3).

In all analyses below, we controlled for age, gender, race, ethnicity, handedness, and evidence of drug consumption on day of testing, which mostly associated with BMI (SI Appendix, section Results, Fig. S2). When presenting and interpreting phenotypic associations, we controlled for family structure to avoid inflated effect sizes and standard errors (e.g., 25). The behavioural genetics analysis did not control for family structure, since this information is needed for modelling heritability. As socio-economic status (SES) is intertwined with cognitive test scores (26), personality (27), and brain morphometry (28), we also present phenotypic associations controlling for SES (education and income) in the supplementary material. All in-text p-values are provided without correcting for multiple comparisons. False discovery rate (FDR) correction was applied when screening for features within cognitive, personality, and brain factors (Fig. 1,2,5).

**Fig. 1.**
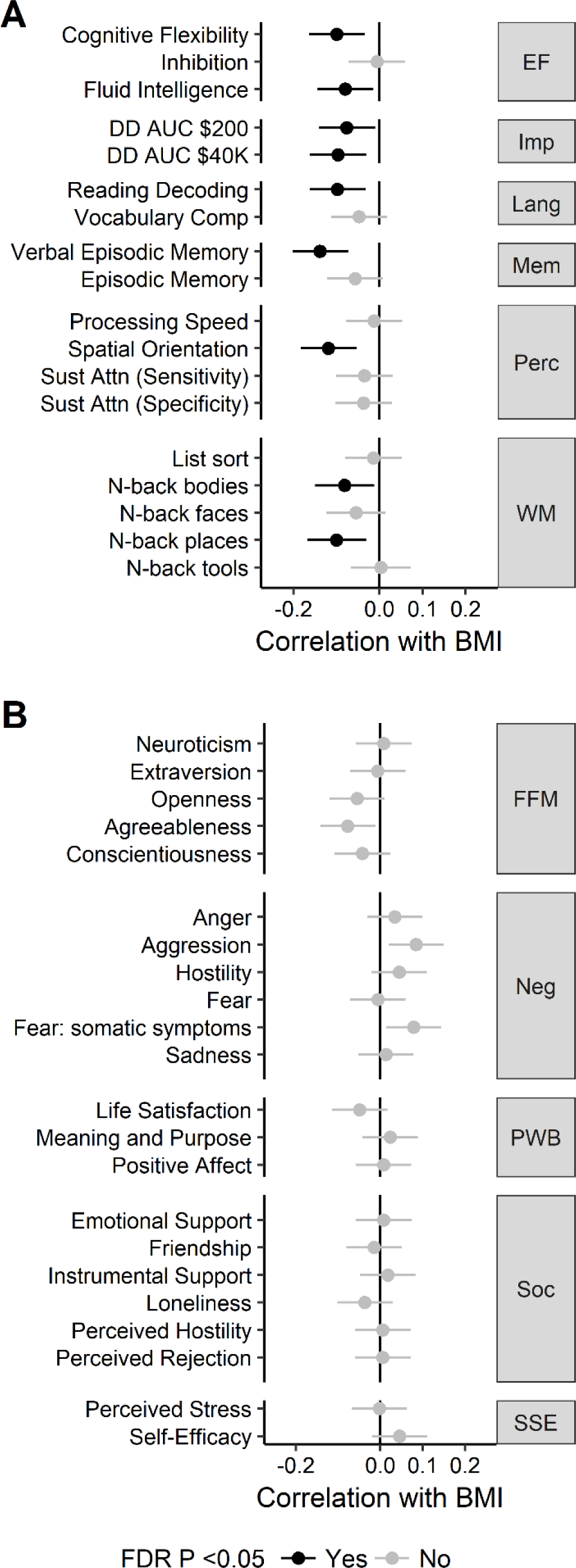
Associations between body mass index (BMI) and (A) cognitive test scores, and (B) personality traits (B). Error bars represent 95% confidence intervals. See Dataset S1, section 1 for explanation of cognitive tests. Numerical values are reported in Dataset S1, section 2. EF=executive function; FFM=Five-Factor Model; FDR=false discovery rate; Imp=(lack of) impulsivity; Lang=language; Mem=memory; Neg=negative affect; Perc=perception; PWB=psychological well-being; Soc=social relationships; SSE=stress and self efficacy; WM=working memory.

**Fig. 2.**
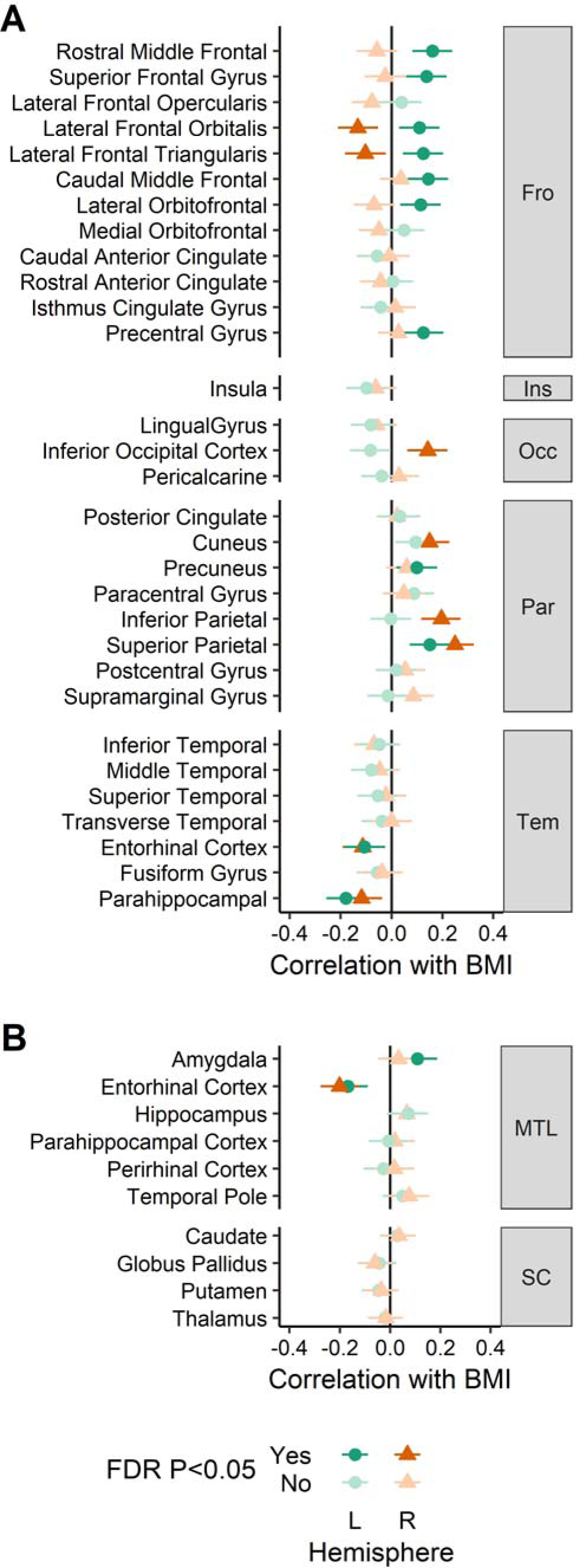
Associations between body mass index (BMI) and brain morphometry. (A) cortical thickness. (B) medial temporal and subcortical regional brain volume. Error bars represent 95% confidence intervals. Numerical values are reported in Dataset S1, section 2. FDR=false discovery rate; Fro=frontal, Ins=insula; L=left; Occ=occipital; Par=parietal; R=right; Tem=temporal; MTL=medial temporal lobe; SC=subcortical.

### Cognitive and Personality Factors

BMI was negatively correlated with the following tests of executive function: cognitive flexibility, fluid intelligence, inability to delay gratification, reading abilities, and working memory. Intriguingly, the strongest effects were present for non-executive tasks measuring visuospatial ability and verbal memory (Fig. 1A). These tasks remained associated with BMI after controlling for SES; controlling for SES reduced the number of executive function tests involved with BMI to cognitive flexibility and inability to delay gratification (SI Appendix Fig. S3A left). No personality test score correlated with BMI when FDR correction was applied (Fig. 1B).

### Brain Morphology

Cortical thickness was estimated from each T1-weighted MRI using CIVET 2.0 software (29). Parcel-based analysis identified negative associations with BMI in right inferior lateral frontal cortex, and bilateral entorhinal-parahippocampal cortex (Fig. 2A & 3A). Positive associations with BMI were found with the left superior frontal cortex, left inferior lateral frontal cortex, and bilateral parietal cortex parcels. Controlling for SES did not change these results (SI Appendix Fig. S4A left). The frontal lobe asymmetry in the BMI association (thinner on the right, thicker on the left) mostly involved the inferior lateral prefrontal areas, such as inferior frontal gyrus. Regional brain volumes were measured for estimation of brain morphology-obesity associations in brain structures not covered by the CIVET cortical thickness algorithm. Medial temporal lobe and subcortical volumes were individually segmented and measured by registering each brain to a labelled atlas using ANIMAL software (30). Volumetric results demonstrated an association between BMI and lower volume of the entorhinal cortex bilaterally, and a positive association of left amygdala volume with BMI (Fig. 2B & 3B). No subcortical region had a significant association with BMI, and results did not change when controlling for SES (SI Appendix Fig. S4B left).

**Fig. 3.**
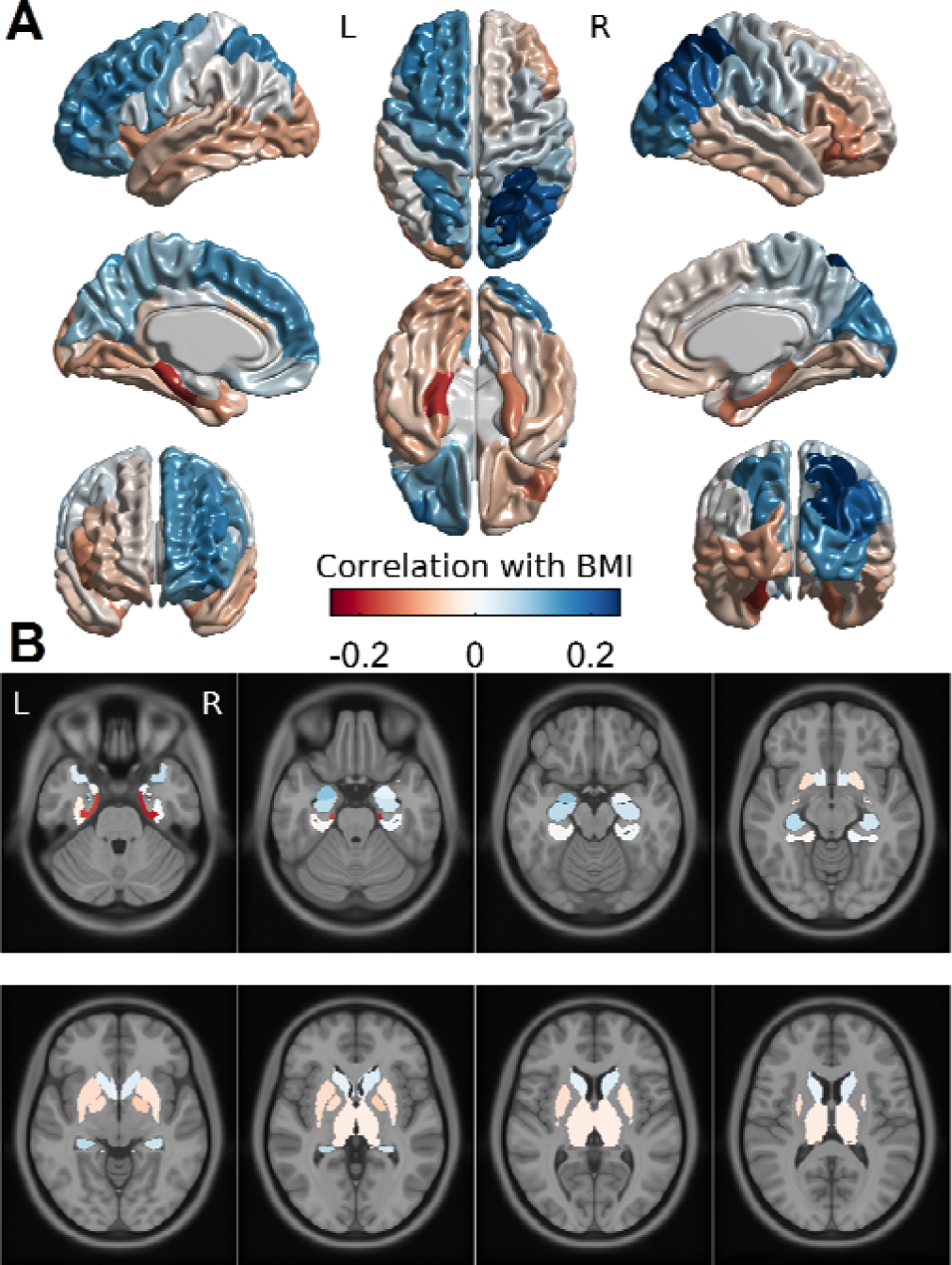
Brain maps of the associations between body mass index (BMI) and (A) cortical thickness and (B) medial temporal and subcortical regional brain volume on a standard brain template in MNI space. Values are the same as in Fig. 2. Colour bar applies to both sub-plots. L=left; R=right.

### Creating poly-phenotype scores

We performed dimension reduction for heritability analyses to reduce measurement noise and avoid multiple testing with redundant measures. Similarly to other recent papers, (20, 27), we used the weights of each individual feature within a neurobehavioural factor (personality test, cognitive test, brain parcel) to create an aggregate BMI risk score or *poly-phenotype score (PPS)*. This is similar to the polygenic score approach in genetics, where the small effects of several polymorphisms are aggregated to yield a total effect score (15, 19, 20, 27). We used the correlation values as weights to multiply each participant’s scaled measurements, and aggregated the results into a single composite variable, the PPS. The PPS reflects the total association of each neurobehavioural factor with BMI. To avoid overfitting, we assigned each 10% of participants the PPS weights obtained from the other 90% (see SI: Data analysis for details).

The associations between BMI and the PPS-s for cognition (correlation with BMI: *r*=0.16, *p*<0.001*, n*=798) and personality (*r*=0.08, *p*=0.017, *n*=888) are slightly higher than the meta-analytic estimates of the pooled association between BMI and cognitive test scores (*r*=0.10, ref: 8) and personality factors (*r*=0.05, ref: 8). BMI had stronger associations with the PPS-s for cortical thickness (*r*=0.26, *p*<0.001, *n*=591), and medial temporal brain volume (*r*=0.23, *p*<0.001, *n*=594). There was no association between BMI and subcortical brain volume (*r*=-0.05, *p*=0.169, *n*=828). To test the generalizability of the PPS approach, we used weights obtained from the full S900 release (SI Appendix Fig. S3 right and S4 right) to test PPS-BMI correlation amongst the unseen additional participants in the S1200 release (referred to as S1200n, n=236). Cortical thickness PPS had essentially unchanged effect size when correlated with BMI in S1200n (SI Appendix, section Results, Fig. S7). At the same time, cognitive and personality PPS-s were less stable (SI Appendix, Section Results, Fig. S7)., likely because the smaller effect sizes of individual features need larger training datasets to reduce inaccuracies, or that the true PPS-BMI effect size was too small to be found just within the S1200n sample.

### Heritability

Bivariate heritability was similarly conducted with the AE model, since the main goal was to explain variance in BMI, for which AE was the best model. All PPS-s were found to be highly heritable, with the A component explaining 36-79% of the variance (Fig. 4A, Dataset S1, section 10). Significant genetic correlations (*r_g_*) were found between BMI and cognitive test scores (*r_g_*=0.25 (*p*=0.002), cortical thickness (*r_g_*=0.45, p<0.001), and medial temporal brain volume (*r_g_*=0.36, *p*<0.001) (Fig. 4B, Dataset S1, section 11). The personality PPS genetic correlation with BMI was not significant (*r_g_*=0.22, *p*=0.052). Molecular evidence relying on linkage disequilibrium score regression has reported effects of similar magnitude between higher cognitive test scores and BMI (rg=-0.22, ref:, 15, rg=-0.18, ref:, 18). Environmental correlations (i.e. correlations between environmental variances) were small and not significant (Dataset S1, section 11). As expected from high heritability of the traits and high genetic correlations, the phenotypic BMI-PPS correlations described in the previous sections were 77-89% explained by genetic factors (Fig. 4C, Dataset S1, section 10).

**Fig. 4.**
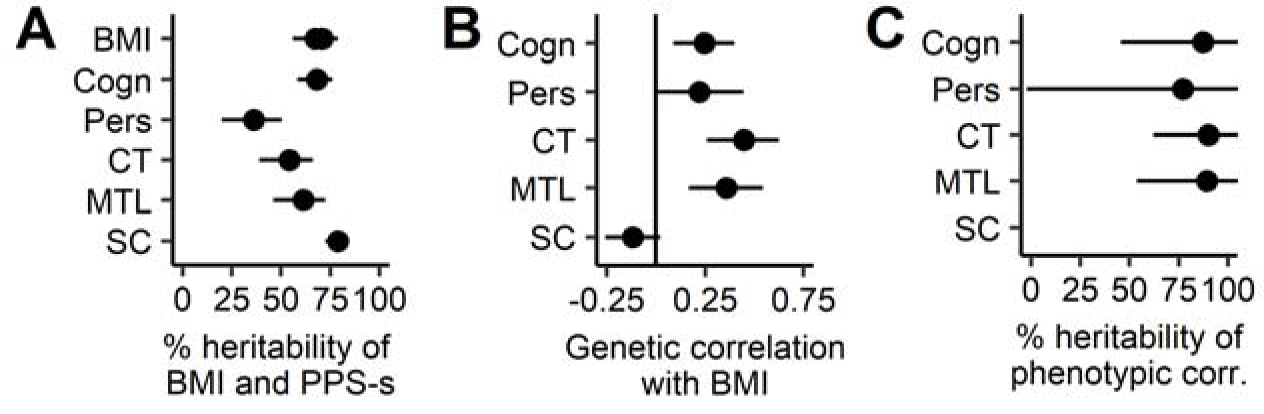
Heritability analysis of the association between poly-phenotype scores (PPS) and body mass index (BMI). (A) Heritability of each trait. BMI has multiple estimates, since it was entered into a bivariate analysis with each PPS separately. (B) Genetic correlations between BMI and each PPS. The genetic correlations are positive, because the PPS-s are designed to positively predict BMI. (C) Heritability of the significant phenotypic correlation between BMI and PPS. Horizontal lines depict 95% confidence intervals. Cogn=PPS of cognitive tests; corr=correlation; CT=PPS of cortical thickness; MTL=PPS of medial temporal lobe volume; Pers=PPS of personality tests; SC=PPS of subcortical structure volumes.

The results broadly replicated when repeating the analysis with just the top features within a PPS, suggesting that PPS based findings summarize the effects of the underlying individual features (SI Appendix Fig. S8). We further replicated the heritability patterns in a separate analysis focused only on the additional participants from the S1200 HCP release (SI Appendix Fig. S9). Additionally, controlling for SES (education and income) did not change the results for brain-based PPS-s. However, the estimates for cognitive test scores and personality became lower and not significant in the S900 release (Figure S10). However, the same estimates were significant in the combined sample S900+S1200, suggesting that the effects of cognition and personality were reduced but not eliminated when controlling for SES.

### Genetic overlap between neurobehavioural factors

Phenotypically, certain PPS-s had small but significant intercorrelations (SI Appendix Fig. S11 upper triangle). After FDR correction, we were able to find two genetic correlations between PPS-s of cognition and cortical thickness (r_g_=0.35), as well as cognition and personality (r_g_=0.33, SI Appendix Fig. S11 lower triangle). Taken together, while the neurobehavioural factors have mostly independent effects on BMI, cognitive test scores may have a small genetic overlap with brain structure and personality.

## Discussion

Cortical thickness, medial temporal lobe volume, and cognitive measures all had covariation with BMI, and their effect on BMI was almost entirely heritable. Similarly, we found genetic correlations between obesity risk scores of cognition, cortical thickness and personality. Together, our results from a large sample support the role of brain and psychological constructs in explaining genetic variance in BMI.

BMI correlated with increased cortical thickness in the left prefrontal cortex and decreased thickness in the right prefrontal cortex, supporting the “right brain” hypothesis for obesity (10). The effect was most prominent in the inferior frontal gyrus (Fig. 2A and 3A). Only preliminary support for the right brain hypothesis has been previously available (13). Right prefrontal cortex has been implicated in inhibitory control (22) and possibly bodily awareness (10). Many neuromodulation interventions (e.g. transcranial magnetic stimulation) aimed at increasing self-regulation capacity often target right prefrontal cortex. On the other hand, effects have also been demonstrated in studies targeting left prefrontal cortex (31).

Cortical thickness results also highlighted the role of temporo-parietal perceptual structures in obesity. Namely, BMI was associated with bilaterally decreased thickness of the parahippocampal and entorhinal cortices, and with mostly right-lateralized increased thickness of parietal and occipital lobes. Volumetric results within the medial temporal lobe supported the role of entorhinal cortex and also suggested that obesity is positively associated with the volume of left amygdala. Emergence of the effects of the right parietal structures together with right prefrontal structures hint at the role of the ventral frontoparietal network, thought to be especially important for detection of behaviourally relevant visual stimuli (32). The parahippocampal and entorhinal cortex are associated with episodic memory and context mediation (33). Similarly, the hippocampus has been associated with the modulation of food cue reactivity by homeostatic and contextual information, and hippocampal dysfunction is postulated to promote weight gain in the western diet environment (34). The amygdala is implicated in emotional and appetitive responses to sensory stimuli, including food cues (35).

Integrating these findings, one could envision a model where obesity is associated with a certain cognitive profile (36). The model starts with a hyperactive visual attention system attributing heightened salience to food stimuli, implicating the ventral visual stream and amygdala. These signals are then less optimally tied into relevant context by the parahippocampal and entorhinal structures, and less well moderated (or filtered) by the prefrontal executive system. This could result in consummatory behaviour driven by the presence of appetitive food signals, which are ubiquitous in our obesogenic environment. An impaired response inhibition and salience attribution model of obesity has been suggested based on the functional neuroimaging literature. Namely, functional MRI studies have consistently identified obesity to associate with heightened salience response to food cues, coupled with reduced activation in prefrontal and executive systems involved in self-regulation and top-down attentional control (e.g., 35). A similar conclusion emerged from a recent resting state network analysis of the HCP data (37), in which obesity was associated with alterations in perceptual networks and decreased activity of default mode and central executive networks.

This brain morphology-derived model has some support from cognitive tests. The role of prefrontal executive control is outlined by our finding of a negative association between BMI and scores on several executive control tasks. Surprisingly, there was no effect of motor inhibition as measured by the Flanker inhibitory task. A relation between obesity and reduced motor inhibition, while often mentioned, has been inconsistent even across meta analyses (7, 8). On the other hand, we found a relationship between decisional impulsivity, measured by delay-discounting, and BMI, replicating previous literature (6, 7, 18). While controlling for education reduced the number of executive tasks associated with BMI, the overall pattern remained the same, suggesting that education level is a proxy for certain executive function abilities.

Intriguingly, BMI was found to be negatively associated with spatial orientation and verbal episodic memory. These tasks tap into the key functions associated with entorhinal and parahippocampal regions implicated in our study (33). Therefore, both cognitive and brain morphology features propose that the increased salience of food stimuli could be facilitated by dysregulated context representation in obesity.

Regarding personality, we were unable to find any questionnaire-specific effects, notably with respect to Neuroticism and Conscientiousness, both often thought to be associated with obesity (5–7). There are potential explanations for this negative finding. First, the meta-analytical association between various personality tests and BMI is small (r=0.05, ref: 7), for which we might have been underpowered after p-value correction. Second, controlling for family structure likely further reduced the effect sizes (25). Third, the personality-obesity associations tend to pertain to more specific facets and nuances than broad personality traits (38), therefore, further analysis with more detailed and eating-specific personality measures is needed in larger samples.

All the associations discussed here were largely due to shared genetic variance between neurobehavioural factors and BMI. This is in accordance with recent molecular genetics evidence that 75% of obesity related genes express preferentially in the brain (4). Similarly, the genetic correlation between cognition and BMI uncovered in our sample is at the same magnitude as molecular estimates of associations between more specific cognitive measures and BMI (15, 18). The current evidence further supports the brain-gene association with obesity vulnerability.

A possible explanation of the genetic correlations is pleiotropy – the existence of a common set of genes that influence variance in both obesity and brain function. It is possible that people with a higher genetic risk for obesity have also genetic propensity for the brain and cognitive patterns outlined here. It is then likely that also interventions could influence both obesity and brain function. For instance, regular exercise can support weight management (39), reduce the heritability of obesity (40), and improve cognitive health (41).

However, our results could also support a causal relationship – that the genetic correlation is due to a persistent effect of heritable brain factors on overeating and hence BMI. For instance, we could hypothesize that the heritable obesity-related cognitive profile promotes overeating when high-calorie food is available. As high-calorie food is abundant and inexpensive, the cognitive risk profile could lead to repeated overeating providing an opportunity for genetic obesity-proneness to express. Such longitudinal environmental effects of a trait need not to be large, they just have to be consistent (42, see discussion in 43). Of course, a reverse scenario is also possible – obesity leads to alterations in cortical morphology due to the consequences of cardiometabolic complications, including low-grade chronic inflammation, hypertension, and vascular disease (reviewed in 9, 44). However, we find this hypothesis less plausible as global brain atrophy due to metabolic syndrome is mostly seen in older participants, whereas the current sample had a mean age of 29. Young adults often experience “healthy or transitional obesity”, where clinical inflammation levels (45) and other cardiometabolic comorbidities have not yet developed (46).

We found neurobehavioural PPS-s to have occasional phenotypic and genetic correlations with each other. Here, it is hard to argue against pleiotropy playing a role. While one could reasonably expect that at least part of the variation in cognitive performance would be shaped by brain morphometry (22), it is also the case that engaging in education leads to improvement in cognitive test scores (26) and might also lead to changes in cortical thickness (47). The small genetic overlap between cognition, cortical thickness and personality can probably be explained by common pleiotropic roots. At the same time, integrating morphometry and cognitive findings is difficult with this dataset.

From a practical point of view, our work suggests that evidence from psychology and neuroscience can be used to design intervention strategies for people with higher genetic risk for obesity. One way would be modifying neurobehavioural factors, e.g. with cognitive training, to improve people’s ability to resist the obesogenic environment (31, 36). Another path could be changing the immediate environment to be less obesogenic (e.g., 48) so that individual differences in neurobehavioural factors would be less likely to manifest. In any case, obesity interventions should not focus solely on energy content, but also acknowledge the certain neurobehavioural profile that obesity is genetically intertwined with.

The current analysis has limitations. Due to the cross-sectional nature of the dataset, causality between neurobehavioural factors and BMI is only suggestive – longitudinal designs would enable better insight into the causal associations between brain morphology, psychological measures, and BMI or weight gain. BMI is a crude proxy for actual eating behaviours or health status. In addition, there were more normal-weight than obese participants. However, the 25% obesity rate in this sample is close to the published obesity rate of the state of Missouri (31.7%) and the US (36.5%, ref:, 49). Also, we expect that BMI itself and the neurobehavioural mechanisms behind it are continuum processes, therefore all variation in the range from normal-weight to obesity is likely helping to uncover underlying associations. While the measurement of cognition and personality was exhaustive, it lacked some common behavioural tasks like the stop-signal task, or common questionnaires measuring self-control, impulsivity, and eating-specific behaviours, that have been previously associated with body weight (5, 6). Particularly, the common eating-specific behaviours such as uncontrolled eating (50) are likely better candidates for explaining brain morphology-BMI associations as they are more directly related to the hypothesized underlying behaviour.

One has to be careful in translating individual differences in cortical thickness in normal populations to underlying neural mechanisms. Diverse biological processes have been suggested to influence MRI-based cortical thickness measures, ranging from synaptic density to apparent thinning due to synaptic pruning and myelination (summarized in 49, 50). A definitive model of the underlying mechanism that links normal variations in cortical thickness to differences in brain function cannot be given, as cortical thickness has not been mapped with both MRI and histology in humans (52). Still, the associations between cortical thickness and BMI in one sample were able to predict BMI in a new separate sample, suggesting that the pattern is robust. Our conceptual interpretation of the significance of cortical thickness patterns has support from measures of both brain structure and cognitive function.

Relying on PPS-s prevented us from analyzing detailed interactions between cortical thickness and cognitive function and their genetic overlap with each other. However, given the relatively small associations between PPS-s, and the number of candidate measures that could be expected to interact with one another, we believe it would have been hard to find an association that would have survived multiple testing correction. Future, focused, hypothesis-driven studies have to further elucidate the neurobehavioural mechanisms behind obesity proneness.

In summary, the current analysis provides comprehensive evidence that the obesity-related differences in brain structure and cognitive tests are largely due to shared genetic factors. Genetic factors also explain occasional overlap between neurobehavioural factors. We hope that increasingly larger longitudinal data sets and dedicated studies will help to outline more specific neurobehavioural mechanisms that confer vulnerability to obesity, and provide a basis for designing informed interventions.

## Methods

Data were provided by the Human Connectome Project (24). Certain people were excluded due to missing data or not fulfilling typical criteria. Exclusion details, demographics and family structure are summarized in SI Methods and Table S1. Software pipelines for obtaining features of cortical thickness and brain volume are described in SI Methods. Analysis scripts to reproduce results presented are available at: osf.io/htx7u.

SI Appendix Fig. S1 provides a schematic pipeline for data analysis. Details of each data analysis step are outlined in SI Methods. We describe how PPS weights are obtained through cross-validation and how the weights generalize to a separate dataset (S1200n). We further describe the main principles of twin and sibling-based heritability analysis and replication of these findings using individual features instead of PPS-s, and replication in a separate dataset (S1200n). Finally, the software and packages used are listed.

## Supplementary Information Text

### Methods

#### Participants

Data were provided by the Human Connectome Project (24) WU-Minn Consortium (Principal Investigators: David Van Essen and Kamil Ugurbil; 1U54MH091657, RRID:SCR_008749) funded by the 16 NIH Institutes and Centers that support the NIH Blueprint for Neuroscience Research; and by the McDonnell Center for Systems Neuroscience at Washington University,

The analyzed data were split between the S900 data release (964 participants) and the S1200 data release (236 additional participants). We treated the S900 as the main analysis sample and results from this sample are reported throughout the paper. At times, we used unique participants from the S1200 release for replication, referred to as S1200n. For the main analysis sample, we applied the following exclusion criteria, as these might confound brain-obesity associations: people with missing values on crucial variables, such as age, BMI, education, income, gender, race, and ethnicity (n=6), hypo/hyper thyroidism (n=4), other endocrine problems (n=16), underweight (BMI <=18, n=9), and women who had recently given birth (n=9). In addition, as we used family information to control for participants’ relatedness, we excluded participants that were half-siblings to other participants (n=31). The same exclusions were applied to S1200n (n=11).

The final main analysis dataset consisted of 895 participants, demographics of which are summarized in Table 1. The sample had good gender balance and variation in BMI and income. As limitations, the sample was relatively young and well educated, and BMI distribution was slightly less obese compared to current prevalence estimates for Missouri or the US as a whole (MO: 31.7%, US: 36.5%, ref:, 49). Most people were white and non-Hispanic, however other races-ethnicities were also represented. The participants were nested into 384 families, typically having 1 to 3 siblings in the dataset. For comparison, we also provide the same statistics for the S1200n sample, as well as a subset of S1200n sample in which no participant is related to the S900 sample.

For the heritability analysis between each neurocognitive factor and BMI, we randomly chose one sibling pair per family, ensuring that the pair had complete data. Non-twin sibling pairs were considered equivalent to dizygotic twin pairs with respect to heritability analyses once data was residualized for age and gender. If multiple sibling pairs within a family had complete data, we prioritized choosing monozygotic twin pairs and dizygotic twin pairs over non-twin sibling pairs. Depending on the neurocognitive factor, the heritability analysis was conducted on 46-111 pairs of monozygotic twins (median=97) and 60-202 pairs of dizygotic twins and siblings (median=176).

#### Measures

##### Psychological measures

Participants completed an extensive set of questionnaires and cognitive tests (see 53, 54 for an overview). In the current analysis, we included 22 questionnaires and 18 cognitive tests (see Fig. 2 and Dataset S1, section 1 for complete list). Here we refer to the set of questionnaire results as personality variables, as personality encompasses various patterns of what people want, say, do, feel, or believe (55). Based on our previous review (6) we chose cognitive tests capturing aspects of executive function, memory, and language.

##### Cortical thickness

All T1-weighted MRI images were processed using the CIVET pipeline (version 2.0) (29, 56, 57). Processing was executed on the Canadian Brain Imaging Network (CBRAIN) High Performance Computing platform for collaborative sharing and distributed processing of large MRI datasets (58). Briefly, native T1-weighted MRI scans were corrected for non-uniformity using the N3 algorithm (59). The corrected volumes were masked and registered into stereotaxic space, and then segmented into gray matter (GM), white matter (WM), cerebrospinal fluid (CSF) and background using a neural net classifier (60). The white matter and gray matter surfaces were extracted using the Constrained Laplacian-based Automated Segmentation with Proximities algorithm (61, 62). The resulting surfaces were resampled to a stereotaxic surface template to provide vertex based measures of cortical thickness (63). All resulting images were visually inspected for motion artefacts by experienced personnel and then subsequently processed through a stringent quality control protocol, which only 641 of the 894 participants in our initial cohort passed. In the S1200n, 144 of the 214 passed. For those participants who passed, cortical thickness was then measured in native space using the linked distance between the two surfaces across 81924 vertices and a 20mm surface smoothing kernel was applied to the data (64). The Desikan–Killiany–Tourville (DKT) atlas was used to parcellate the surface into 64 cortical regions (65). Cortical thickness was averaged over all vertices in each region of interest for each subject (66) and the effect of mean cortical thickness was regressed to allow for regional analysis (67). After participant exclusions, data was available for 591/137 participants in the S900/S1200n samples.

##### Volumetric estimates

Because the CIVET cortical thickness method does not cover all medial temporal and subcortical structures, we used volumetric estimates for these brain regions. For subcortical volumetric estimation, T1-weighted scans of the subjects were pre-processed through a computerized pipeline (n=899). Image denoising (68), intensity non-uniformity correction (59), and image intensity normalization into range (0-100) using histogram matching were performed. After preprocessing, all images were first linearly (using a 9-parameter rigid registration) and then nonlinearly registered to an average template (MNI ICBM152) as part of the ANIMAL software (30, 69). The subcortical structures, i.e., thalamus, putamen, caudate, and globus pallidus were segmented using ANIMAL by warping segmentations from ICBM152 back to each subject using the obtained nonlinear transformations. The medial temporal lobe structures, i.e. hippocampus, amygdala, temporal pole, and parahippocampal, entorhinal and perirhinal cortices, were segmented using an automated patch-based label-fusion technique (70). The method selects the most similar templates from a library of labelled MRI template images, and combines them with a majority voting scheme to assign the highest weighted label to every voxel to generate a discrete segmentation. Quality control was performed on the individual registered images as well as the automated structure segmentations by visual inspection, and inaccurate results were discarded. In S900, 648 participants passed the quality control for medial temporal lobe structures, ad 895 for subcortical structures. Within S1200n, of the 214 participants, 212 passed the quality control for subcortical structures, and 174 passed the quality control for medial temporal lobe. After exclusions, the S900/S1200n samples included data from n=828/204, 8 parcels per subjects for the subcortical structures, and n=594/166, 12 parcels for the medial temporal lobe structures.

#### Data Analysis

##### Analyzing each feature

A schematic pipeline of the analysis is displayed in SI Appendix Fig. S1. Data from all neurocognitive factors were first residualized for control variables (age, ethnicity, gender, handedness, race) using multiple linear regression. When presenting phenotypic associations, we used a linear mixed model, adding a random intercept for family (SI Appendix Fig. S1), and also varied the involvement of income and education. As BMI was skewed (long-tail at the upper end of the scale), it was log-transformed to achieve a normal-like distribution. Handedness was also log normalized.

For each factor category (cognition, personality, cortical thickness, medial temporal volume, subcortical volume), factor-BMI relationships were assessed using univariate correlation between each brain parcel or test score and BMI. We initially also tried using a partial least squares (PLS) correlation approach, which is a multivariate technique suited to handling correlated predictors (71, 72). However, the PLS estimates were extremely close to univariate correlations, therefore univariate correlations were preferred for simplicity. As a result, we received an estimate of the relative contribution (weight) of each predictor within a given factor. Estimates used in this study are presented in Dataset S1, section 2.

##### Creating poly-phenotype scores

To summarize effects for each neurocognitive factor, we created an aggregate BMI risk score or *poly-phenotype score (PPS)* for each neurocognitive factor. This was inspired by the polygenic risk score approach, where the effects of single-nucleotide polymorphisms are added up to form a total genetic score (73). Specifically, we used the correlation-derived weights to multiply each participant’s measured values, and aggregated the results into a single composite variable for a given factor, the PPS. A PPS would reflect the total association that a given factor has with BMI. Even though only some features within a neurobehavioural factor had significant effects on BMI, and certain features correlated with each other (see Datasets S3-S7), both our testing (see SI Results) and recommendations by others (74) lead us to not apply p-value cutoffs, clumping, or pruning, as excluding these steps does not hurt predictive ability and improves transparency (74). PPS-s have a mean of 0 but varying standard deviation, depending on the number of features and their effect sizes (Dataset S1, section 8).

We used cross-validation principles to avoid and test for overfitting. Namely, we divided participants into 10% folds. Each 10% fold received the correlation weights from the remaining 90% of the sample. As the result, we received one PPS vector for each factor, where each participant’s score was based on out-of-sample prediction. When creating the 10% folds, we created folds for each factor separately, as each factor has a different number of available data points, ensuring that folds were as equal in size as possible. We also ensured that siblings from the same family were in the same fold. Therefore, no data from family members were used in calculating both the correlation weights and performing out of sample predictions.

To test the robustness of PPS-s, we first tested the impact of not pruning and applying p-value cutoffs. In a pruned PPS, features are omitted that a) correlate above criterion to another feature and b) have lower correlation with BMI than the other feature (75). In a PPS with p-value cut-off, features are omitted that have an above-criterion uncorrected p-value when correlated with BMI Neither pruning nor a p-value cutoff improved the predictive ability of the PPS-s (see SI Results).

We further tested the predictive ability of PPS scores by applying the weights created on the full S900 release to predict BMI in the S1200n release (new participants only), which we did not touch before predicting. As 101 participants within the S1200n were related to participants in the S900, we also tested the predictive ability in the subset of S1200 that was not related to S900 (n=124).

##### Heritability analysis

In the heritability analysis, a typical behavioural genetics decomposition uses relatedness assumptions between individuals to divide variance in a trait to the following components: genetic variance (A, additive and interactive effects), shared environmental variance (C, family and shared school effects), and unique environmental variance (E, unique experience and measurement error). The assumptions are: 100% of genetic variance shared between monozygotic twins, 50% of genetic variance shared between dizygotic twins and sex-and gender residualized siblings, 100% of family environment shared by all siblings, 0% unique variance shared between siblings. Such decomposition is called univariate heritability.

Besides establishing univariate heritability, one can also conduct heritability analysis on the covariance between two traits. For instance, a genetic correlation is the correlation between the A components of trait 1 and trait 2. A bivariate heritability analysis decomposes the phenotypic correlation between trait 1 and trait 2 into A, C, and E components.

Heritability analysis was conducted on PPS scores not residualized for family structure, as this information is used in heritability modelling. We then ran bivariate heritability analyses separately between each PPS and BMI, which provided univariate heritability estimates of the PPS-s and BMI, genetic and environmental correlations between the univariate estimates of PPS-s and BMI, and bivariate decomposition of the phenotypic correlation between each PPS and BMI. We used the AE model, since BMI was best explained by an AE model, as opposed to an ACE model, based on Akaike Information Criterion (AIC) (Dataset S1, section 9). Similar AIC patterns were present for bivariate models (SI Appendix Fig. S12, Dataset S1, section 12). We report only standardized A estimates in the main results, as in the univariate and bivariate analysis of the AE model, E=100-A. Also, no environmental correlations were significant. All standardized and unstandardized estimates are reported in the supplementary materials (Datasets S10-S11).

##### Analysis software

Analysis was conducted in Microsoft R Open 3.4.0 (76), using May 2017 version of packages abind, car, caret, cowplot, corrplot, ggplot2, lme4, MuMIn, pbkrtest, plyr, psych, synthpop, tidyr, WriteXLS (77–92). Cortical thickness was plotted using Surfstat (93) in MATLAB (94). Heritability analysis was conducted using OpenMX (95), adapting scripts provided by the Colorado International Twin Workshop (96).

### SI Results

#### Control variables

Age, gender and race related to BMI, demonstrating the need for residualizing (SI Appendix Fig. S2). Marginal R^2^ explaining only fixed effects was 0.07, and conditional R^2^ explaining both fixed and random effects was 0.38, highlighting the effect of family structure. When controlling for education and income, education was a significant additional predictor, with total model R^2^ being 0.09 and conditional R^2^ 0.37. Further, controlling for family structure in a nested model as random intercept improved model fit (AIC dropped from 7006 to 6895 / 6978 to 6885 when controlling for education and income), suggesting that family nesting needs to be taken into account.

#### Robustness of PPS-s

Similarly to genetic literature (74), we found that pruning features or applying a p-value threshold does not change the predictive ability of the PPS-s (SI Appendix Fig. S5 & S6).

To test the generalizability of the PPS approach, we used weights obtained from the full S900 release (SI Appendix Fig. S3 right and S4 right) to predict the BMI of new participants in the S1200 release (S1200n, n=236), which were not used in any of the initial assessments. As certain participants in the S1200n release were related to participants in the S900, we also tested the PPS performance when they were excluded. As can be seen in SI Appendix Fig. S7, cortical thickness estimates are very similar, no matter the training or testing dataset. Cognition PPS effect sizes were similar to each other, but did not reach statistical significance in the replication sample (S1200n). Personality PPS had unexpectedly high correlation with BMI in the new data. Further research is needed to determine if such effect sizes would further replicate. Medial temporal lobe PPS-s also did not replicate.

#### Heritability replication

We tested whether the PPS-based bivariate analysis patterns would replicate in the S900 dataset, but using unaggregated top individual features within the PPS-s. We chose the 5 individual features from the top predictors of cognition and cortical thickness. As shown in SI Appendix Fig. S8, the individual tasks are comparable with the PPS-s in terms of univariate heritability, genetic correlations, and heritability of phenotypic correlation. However, with genetic correlations, the estimates are non-significant (SI Appendix Fig. S8 B1&B2), suggesting that we are not powered to establish significance of the smaller correlations. Further, the standardized estimates for heritability of the phenotypic correlations (SI Appendix Fig. S8 C1&C2) are noisier and the estimator often failed at estimating standardized confidence intervals. Such failures at individual feature levels highlight the value of PPS-s, which provide more stable estimates at these sample sizes.

We further used participants only in the S1200n release to replicate the bivariate heritability analysis results in new data. PPS weights were obtained from the S900 release. We focused only on participants who did not have siblings in the S900 release. Granted, the power is low because of fewer complete twin pairs available (29 MZ pairs and 30 DZ pairs). The univariate estimate for BMI heritability was [A=64% [95% CI: 41%;79%]. In the bivariate analysis, we were also able to replicate the patterns seen in the main dataset (SI Appendix Fig. S9), however the confidence intervals were often covering 0 or not estimated, likely due to small sample size.

**Fig. S1.**
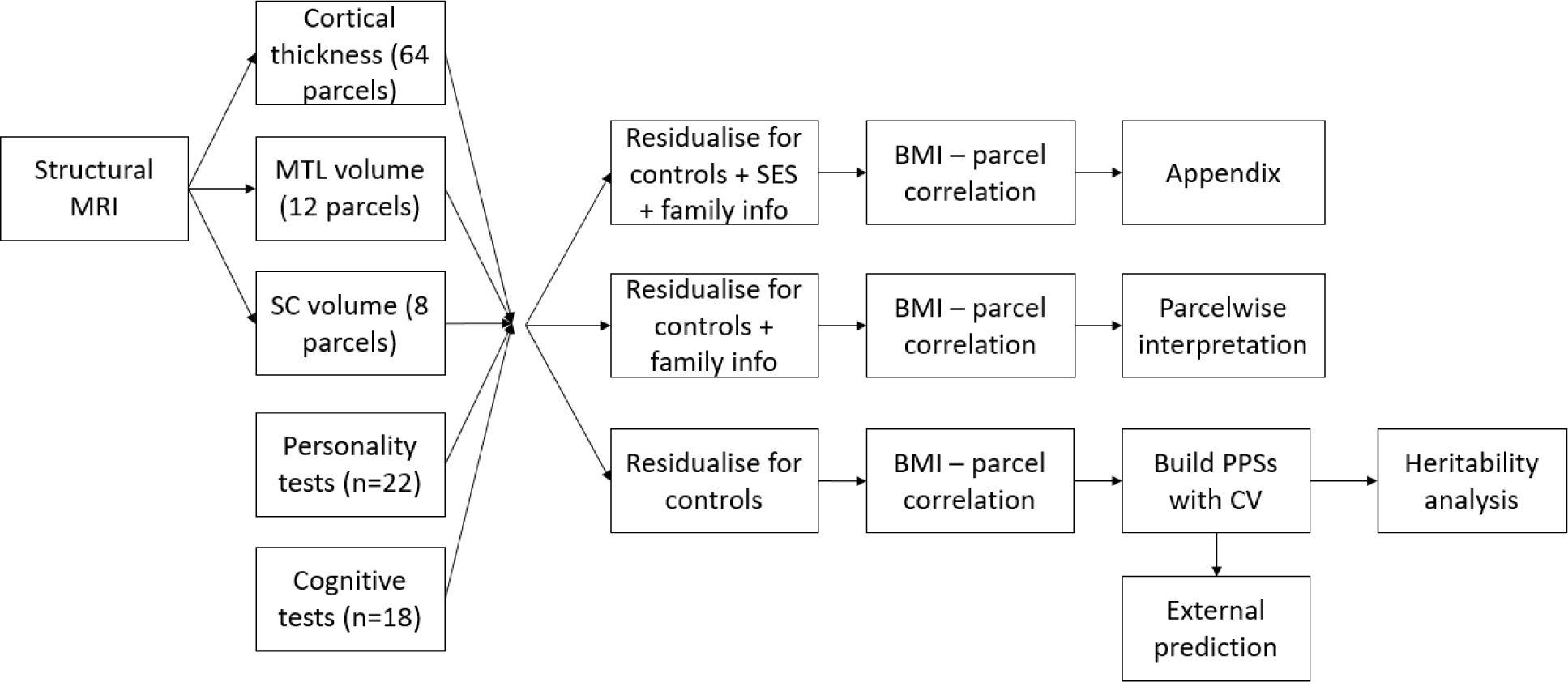
A schematic diagram of the analysis pipeline. All steps were conducted on all neurocognitive factors separately. BMI=body mass index; CV=cross-validation; MTL=medial temporal lobe; MRI=magnet resonance image; PPS=poly-phenotype score; SC=subcortical; SES=socio-economic status (education and income).

**Fig. S2.**
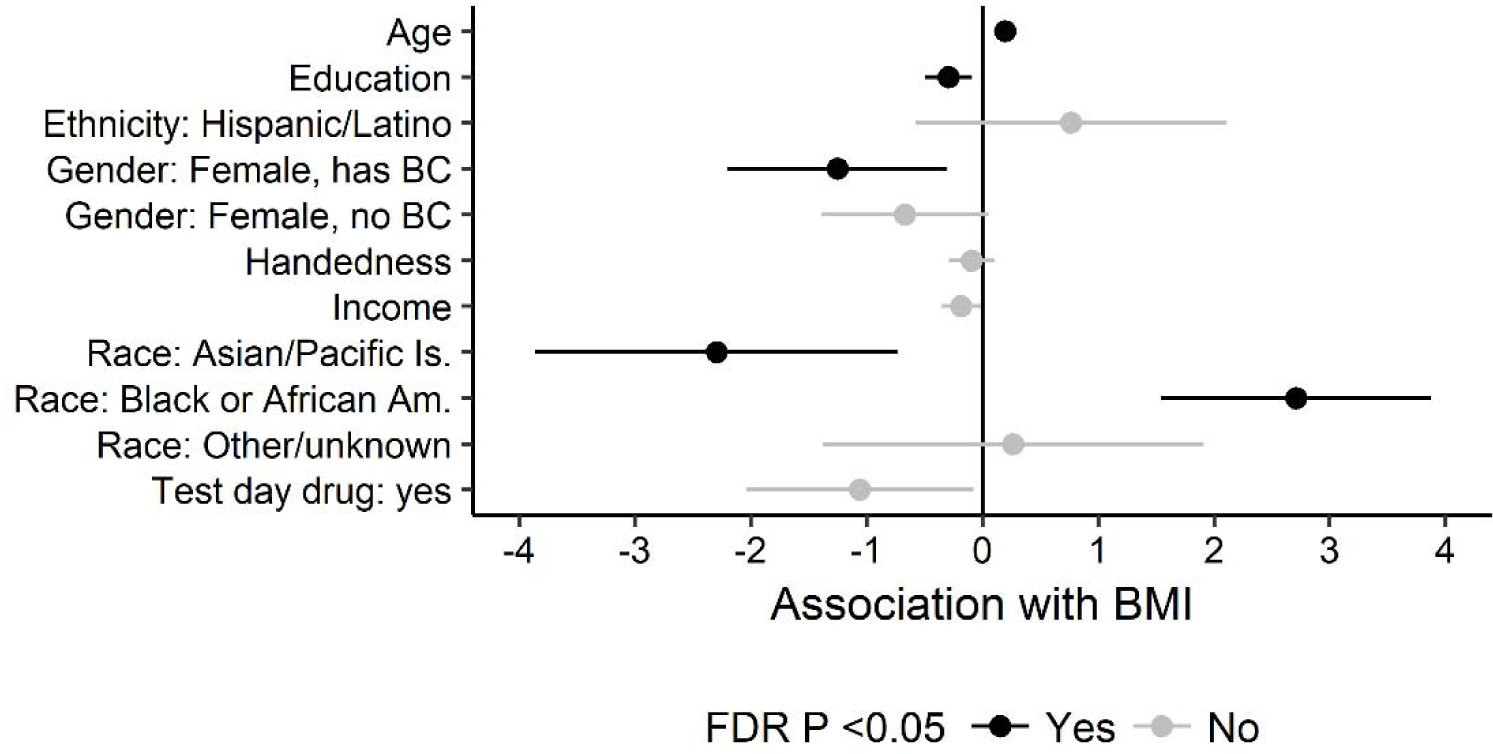
Regression weights of a multilevel linear model nested for family. Lines mark standard 95% confidence intervals. Intercept is 27.37 (standard error: 2.16). For interpretability, regular BMI is unscaled here. Reference groups: Gender: male, Race: white, Ethnicity: not Hispanic/unknown. Am.=American; BC=birth control; Is.=Islander

**Fig. S3.**
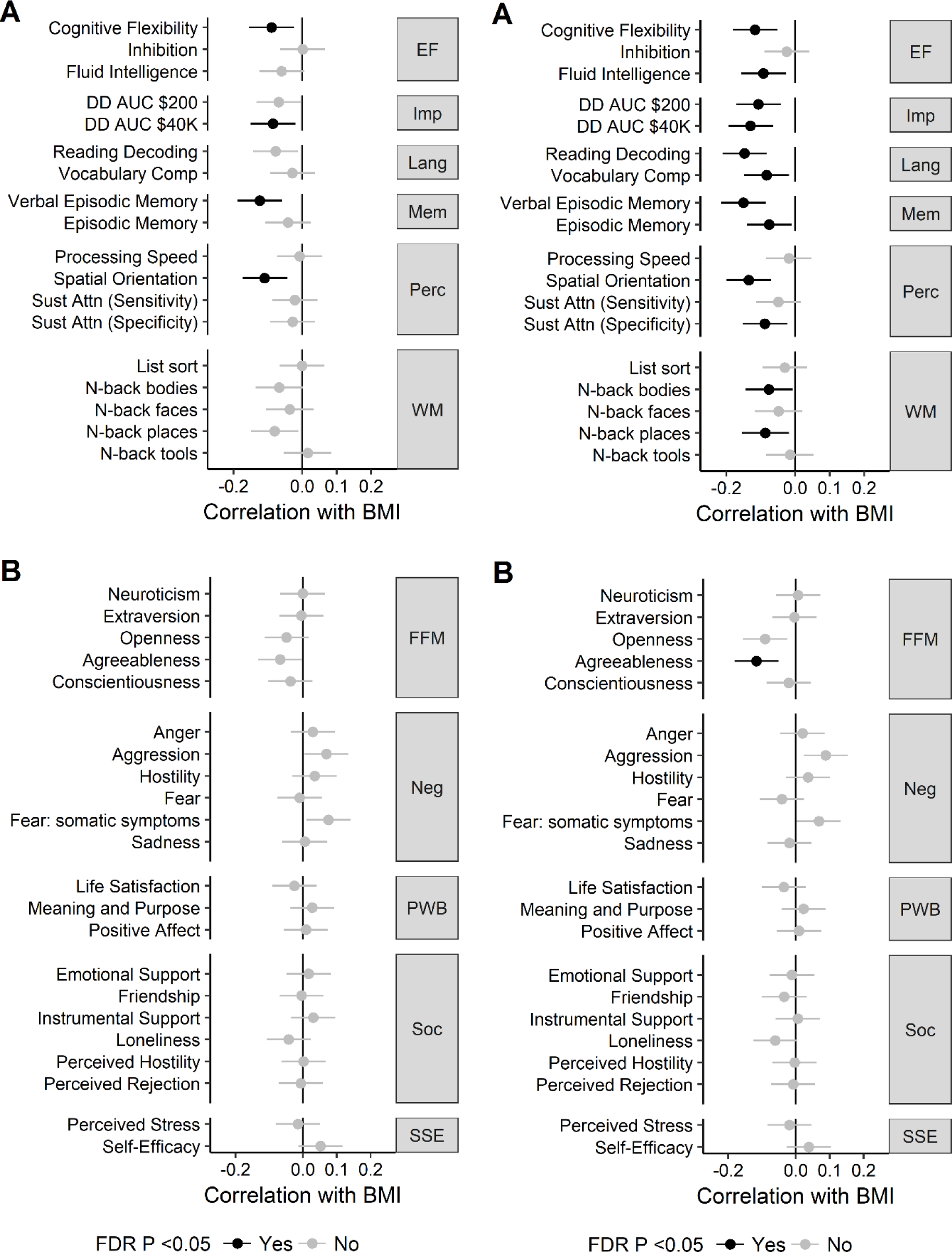
Associations between body mass index (BMI), cognitive test scores (A), and personality traits (B), either when controlling for education, income, and family structure (left), or not controlling for these variables (right). Error bars mark 95% confidence intervals. See Dataset S1, section 1 for explanation of cognitive test names. Numerical values are reported in Dataset S1, section 2. EF=executive function; FFM=Five-Factor Model; FDR=false discovery rate; Imp=(lack of) impulsivity; Lang=language; Mem=memory; Neg=negative affect; Perc=perception; PWB=psychological well-being; Soc=social relationships; SSE=stress and self efficacy; WM=working memory

**Fig. S4.**
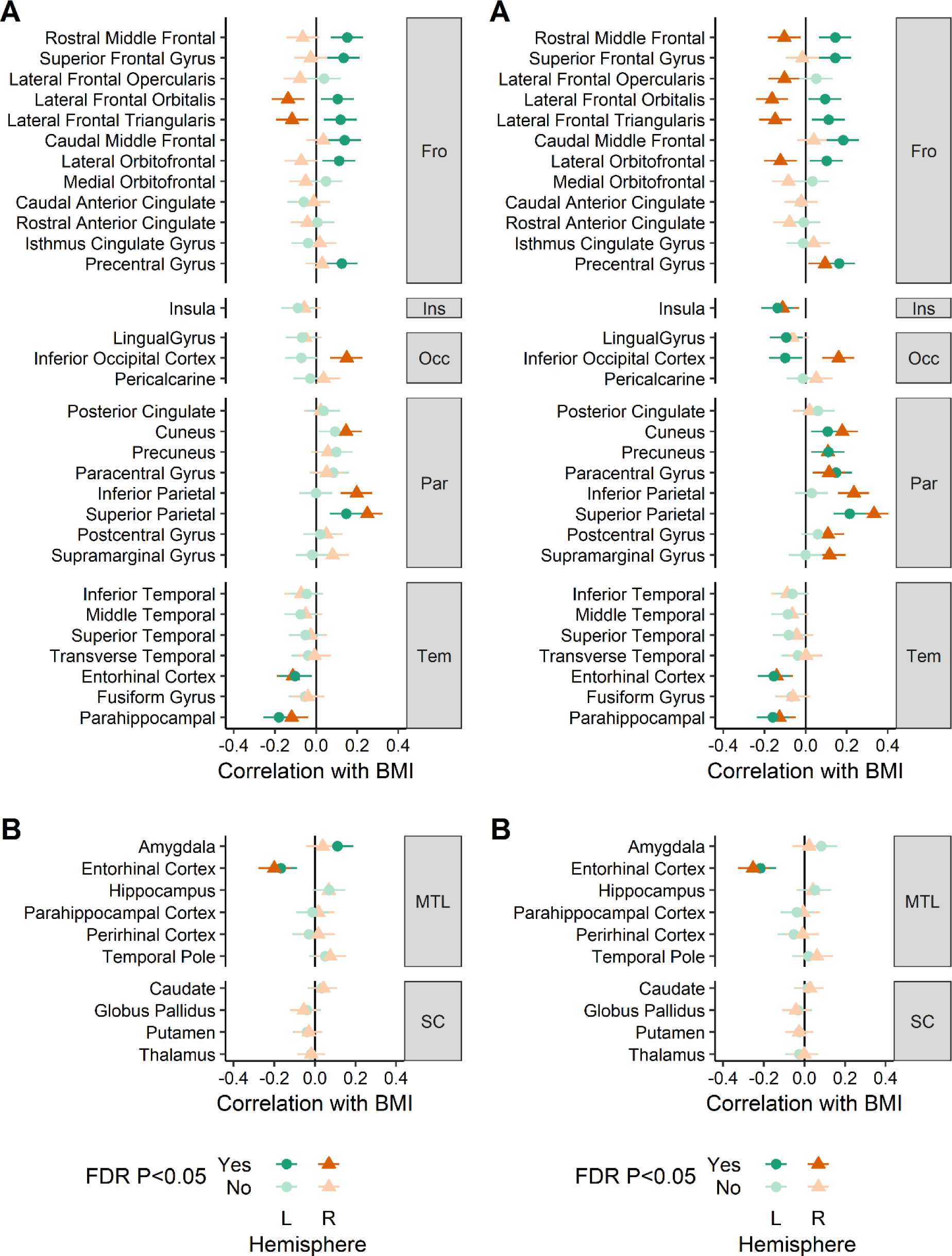
Associations between body mass index (BMI), cortical thickness (A) and regional brain volume (B), either when controlling for education, income, and family structure (left), or not controlling for these variables (right). Error bars mark 95% confidence intervals. Numerical values are reported in Dataset S1, section 2. FDR=false discovery rate; Fro=frontal, Ins=insula; L=left; Occ=occipital; Par=parietal; R=right; Tem=temporal;

**Fig. S5.**
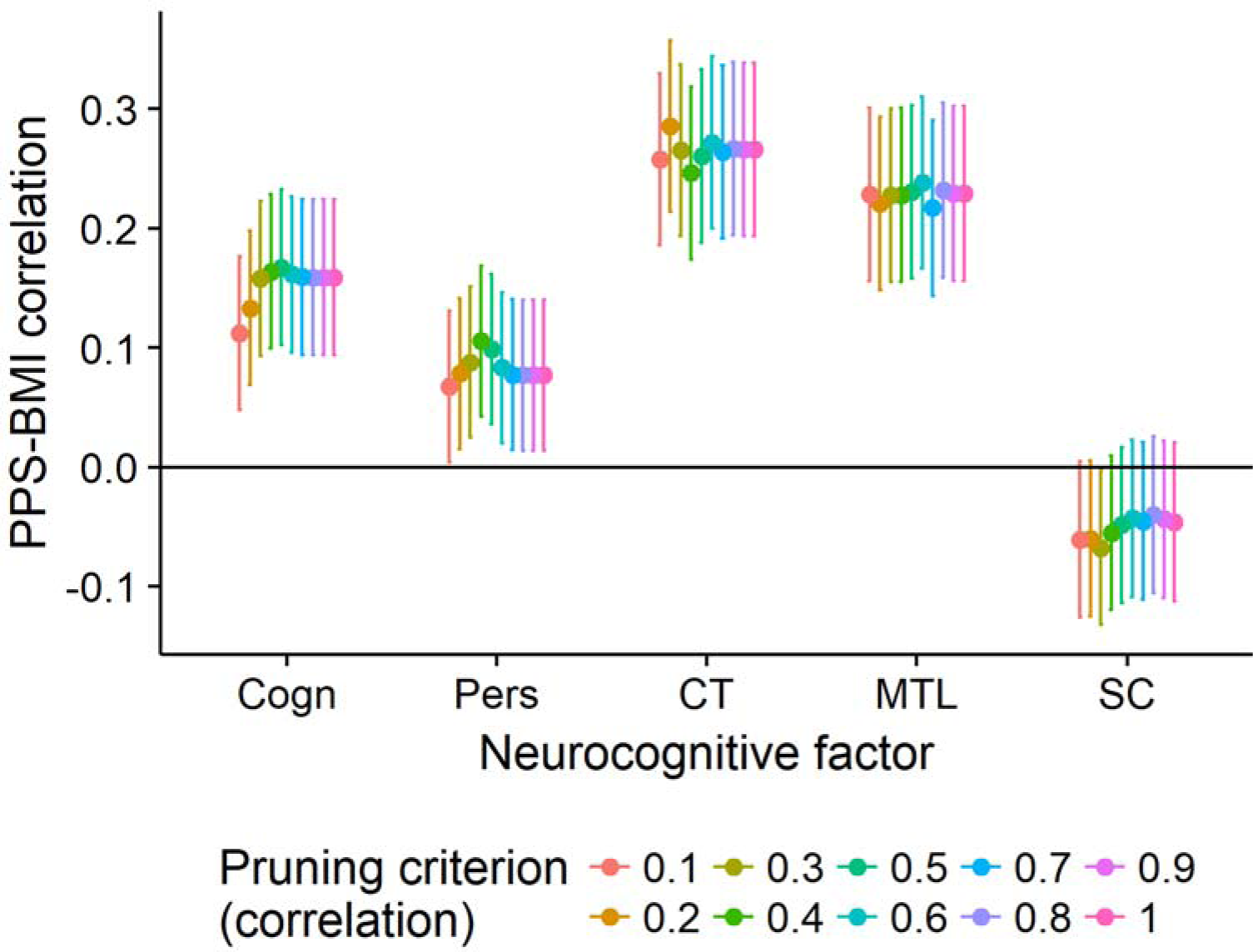
Low impact of pruning to the poly-phenotype scores’ (PPS) associations with BMI. PPS-s were trained and tested within the Human Connectome Project’s S900 release, using cross-validation. Pruning means excluding features that have a higher correlation than set criterion with another feature that associates with BMI. A pruning criterion equal to 1 means no pruning was done. Cogn=PPS of cognitive tests; CT=PPS of cortical thickness; MTL=PPS of medial temporal lobe volume; Pers=PPS of personality tests.

**Fig. S6.**
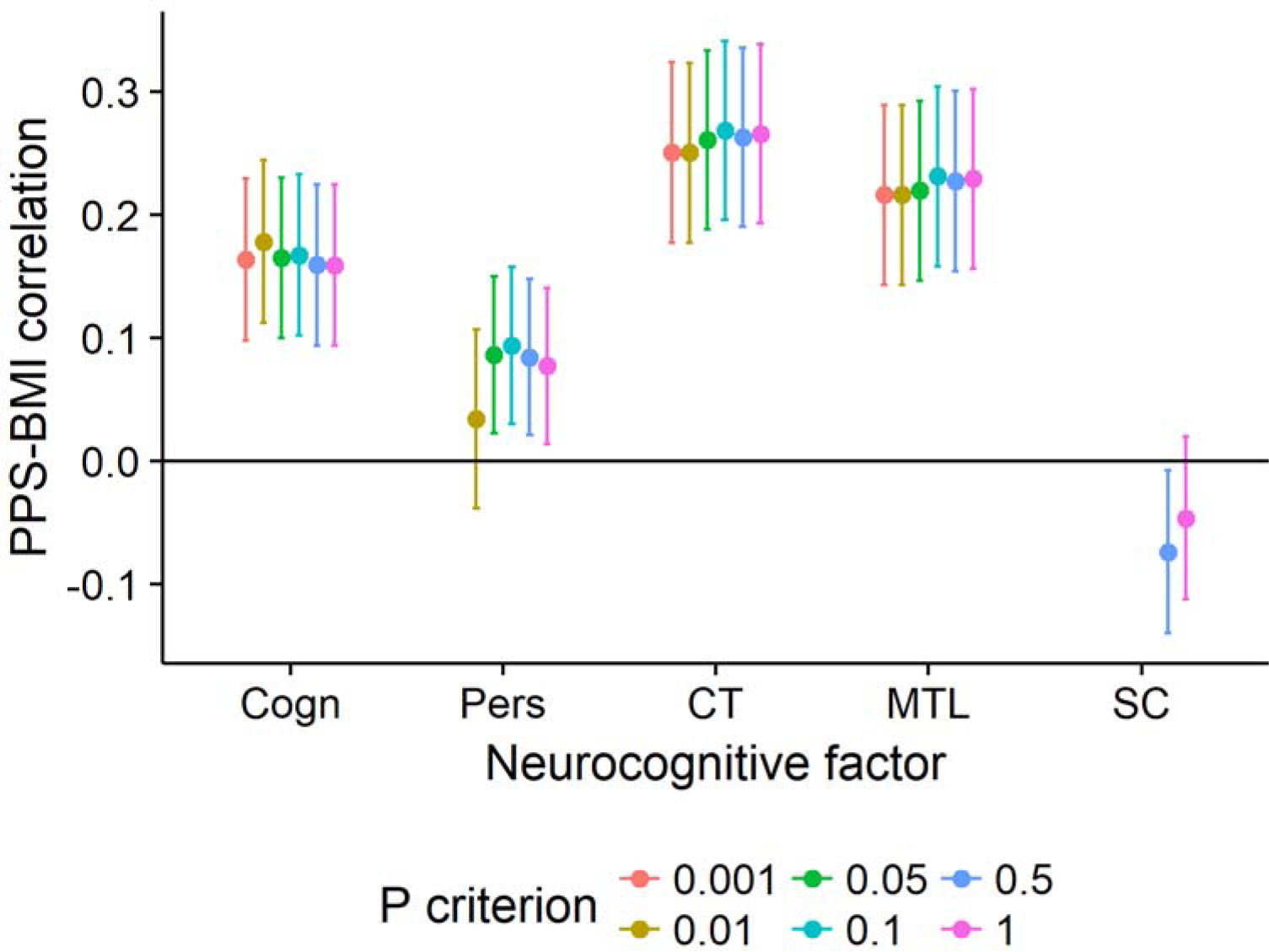
Low impact of excluding features by p value to the poly-phenotype scores’ (PPS) associations with BMI. PPS-s were trained and tested within the Human Connectome Project’s S900 release, using cross-validation. Features with a p value higher than criterion were excluded from the PPS. A p criterion of 1 means no exclusion was done. Cogn=PPS of cognitive tests; CT=PPS of cortical thickness; MTL=PPS of medial temporal lobe volume; Pers=PPS of personality tests.

**Fig. S7.**
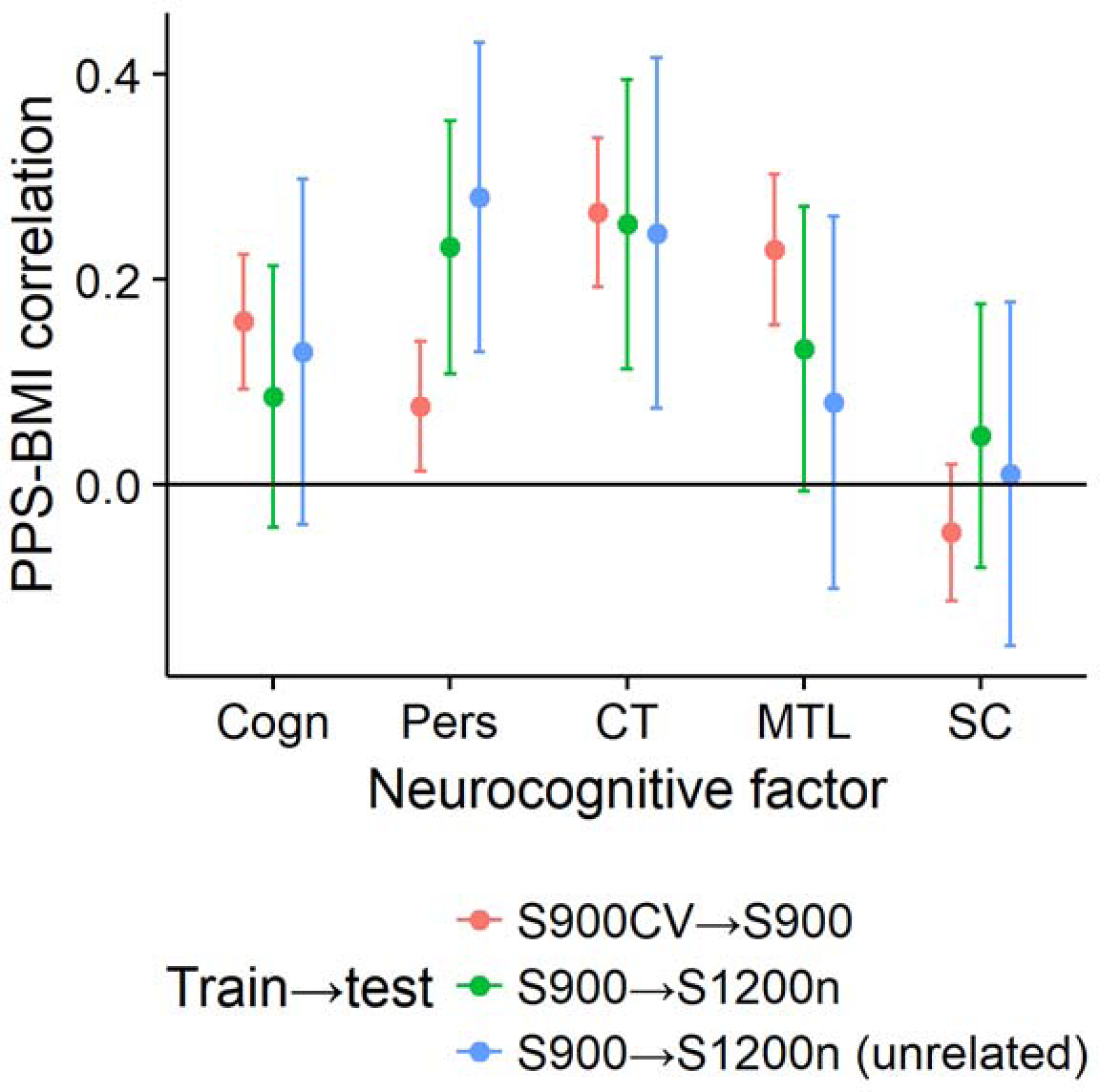
Comparison of poly-phenotype scores’ (PPS) performance in correlating with BMI, depending on training data and test data. S900CV→S900: PPS-s within S900 release trained and tested with cross-validation to avoid bias. These PPS-s are used in heritability analysis. S900→S1200n: PPS-s trained on S900 and tested in full S1200n sample. S900→S1200n (unrelated): PPS-s trained on S900 and tested in S1200n sample not related to S900. Cogn=PPS of cognitive tests; CT=PPS of cortical thickness; CV=cross-validated; MTL=PPS of medial temporal lobe volume; Pers=PPS of personality tests; S900 – Participants in Human Connectome Project’s S900 release; S1200n – participants only in the S1200 release; SC=PPS of subcortical structure volumes;

**Fig. S8.**
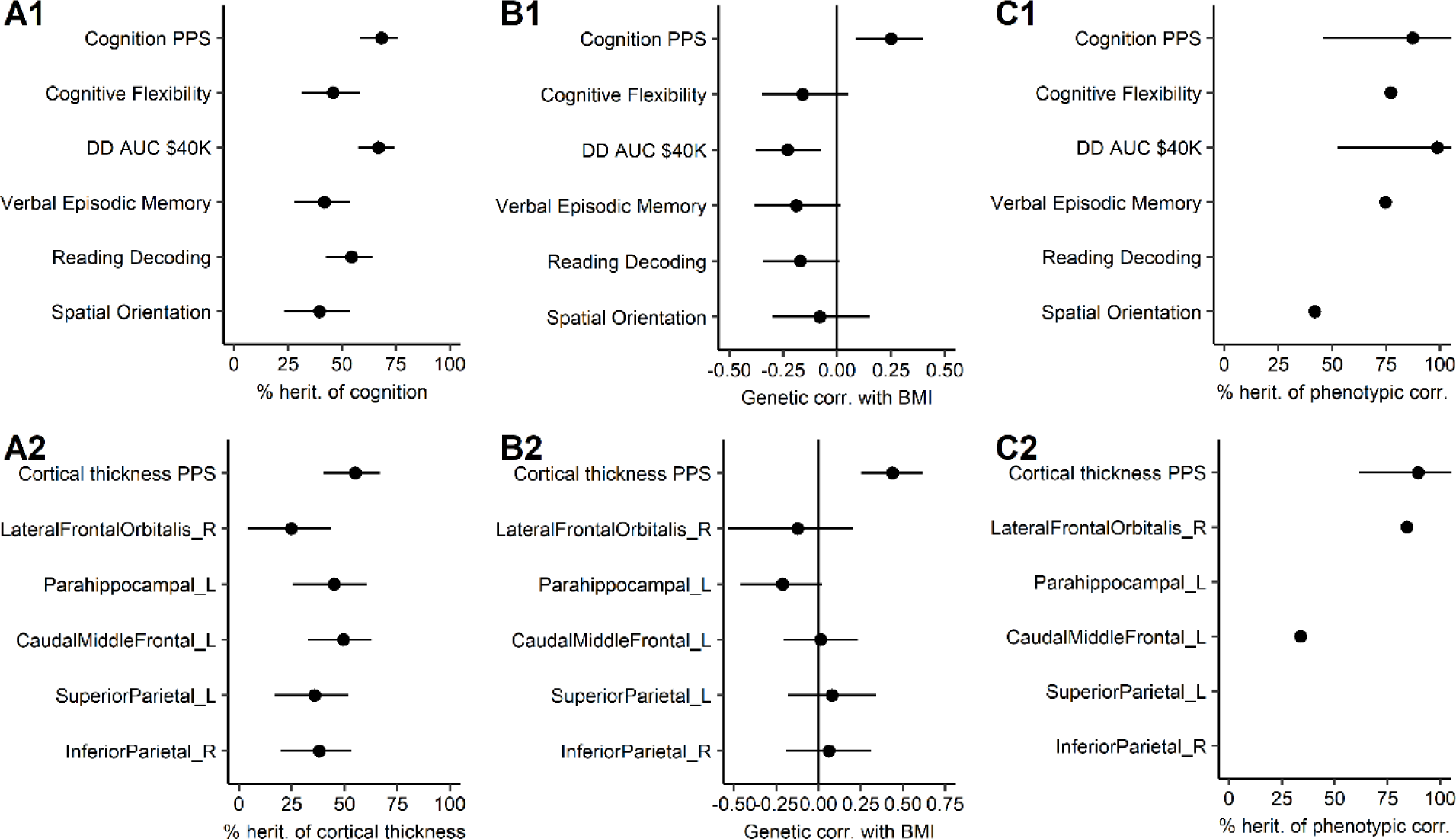
Heritability analysis of the association between poly-phenotype scores (PPS) of cognitive test scores (A1-C1) and cortical thickness (A2-C2), compared with most significant individual features of each PPS. (A) Heritability of each trait. The effect of unique environment (E) is not shown, since E=100-A. (B) Genetic correlations between BMI and each PPS or between BMI and each feature. The PPS-based genetic correlations are positive, because the PPS-s are designed to positively predict BMI. However, individual features can have negative genetic correlations. (C) Heritability of the phenotypic correlation between BMI and PPS or between BMI and each feature. Horizontal lines depict 95% confidence intervals. The estimator failed at estimating certain features. Corr=correlation; L=Left hemisphere; herit=heritability; R=right hemisphere.

**Fig. S9.**
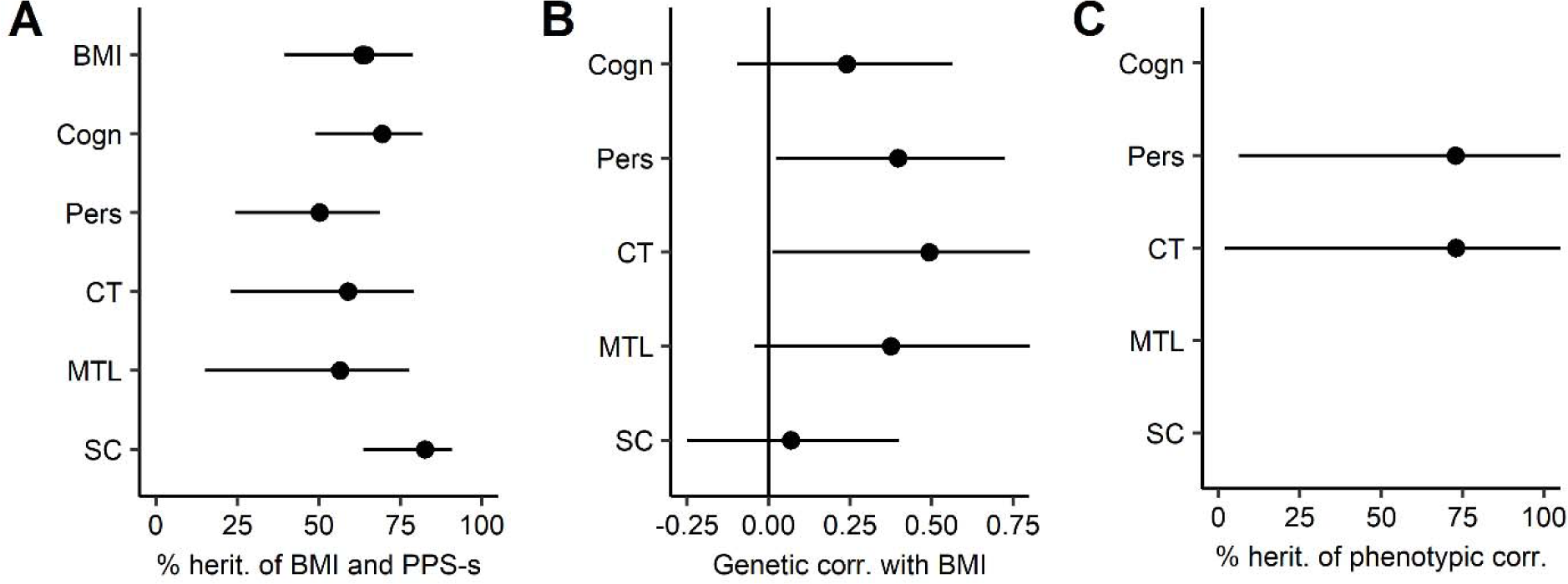
Heritability analysis of the association between poly-phenotype scores (PPS) and body mass index (BMI) in the S1200n sample unrelated to S900. (A) Heritability of each trait. BMI has multiple estimates, since it was entered into a bivariate analysis with each PPS separately. The effect of unique environment (E) is not shown, since E=100-A. (B) Genetic correlations between BMI and each PPS. The genetic correlations are positive, because the PPS-s are designed to positively predict BMI. None of the environmental correlations were significant and therefore not shown. (C) Heritability of the phenotypic correlation between BMI and PPS. Horizontal lines depict 95% confidence intervals. Estimates not shown for PPS-s that did not have significant phenotypic association with BMI. Cogn=PPS of cognitive tests; corr=correlation; CT=PPS of cortical thickness; herit=heritability; MTL=PPS of medial temporal lobe volume; Pers=PPS of personality tests; SC=PPS of subcortical structure volumes

**Fig S10.**
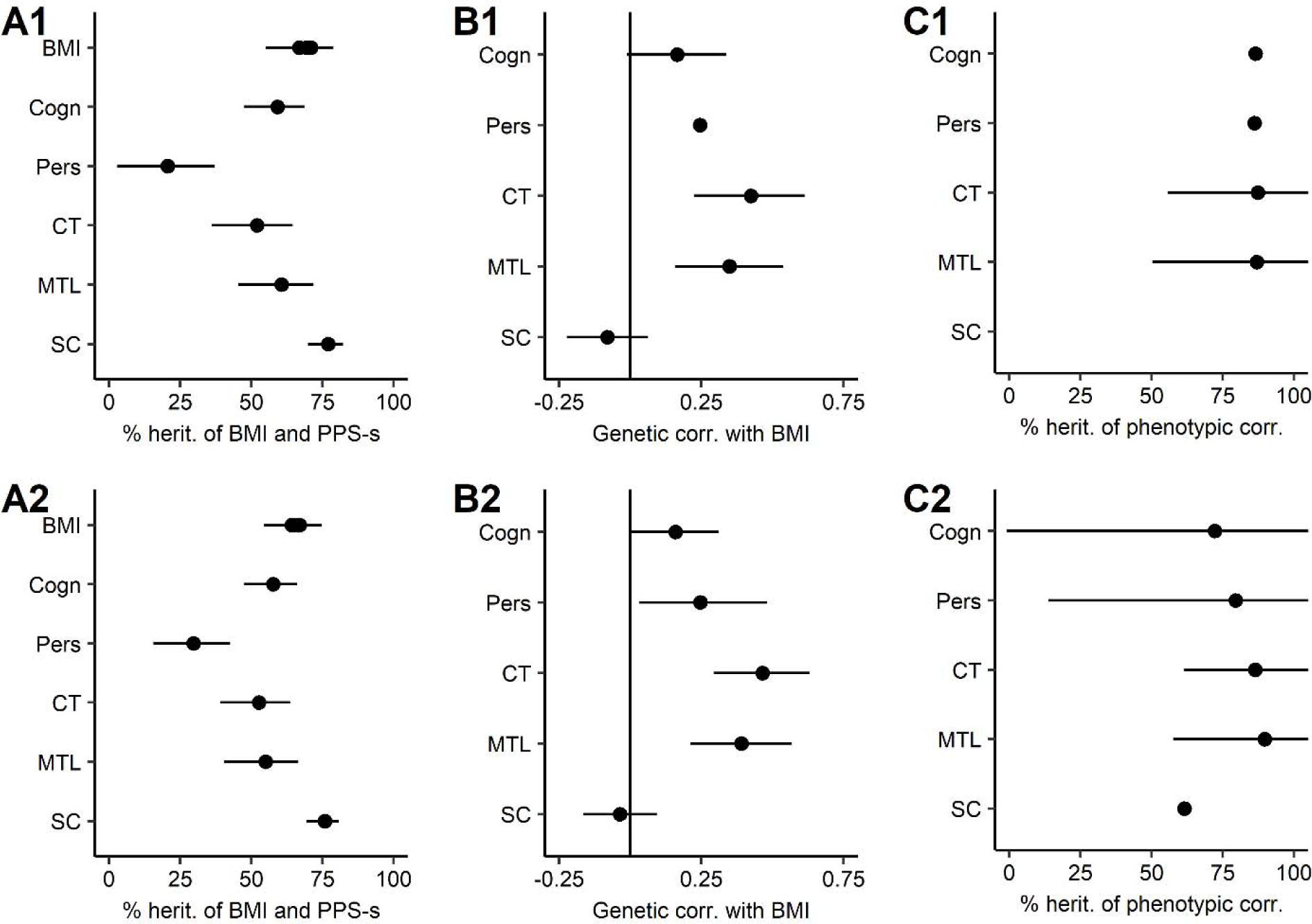
Heritability analysis of the association between poly-phenotype scores (PPS) and body mass index (BMI), when controlling for education and income within the S900 sample (top panel) and in the S1200 sample, where S1200n is added to the S900 sample (bottom panel). As in previous analyses, the PPS weights of S1200n sample are based on S900 sample, S1200n sample just adds statistical power to the S900 based findings. Depending on the neurocognitive factor, the heritability analysis in the combined sample was conducted on 59-135 pairs of monozygotic twins (median=108.5) and 85-259 pairs of dizygotic twins and siblings (median=179). (A) Heritability of each trait. BMI has multiple estimates since it was entered into a bivariate analysis with each PPS separately. (B) Genetic correlations between BMI and each PPS. The genetic correlations are positive, because the PPS-s are designed to positively predict BMI. (C) Heritability of the significant phenotypic correlation between BMI and PPS. Horizontal lines depict 95% confidence intervals. Cogn=PPS of cognitive tests; corr=correlation; CT=PPS of cortical thickness; MTL=PPS of medial temporal lobe volume; Pers=PPS of personality tests; SC=PPS of subcortical structure volumes.

**Fig. S11.**
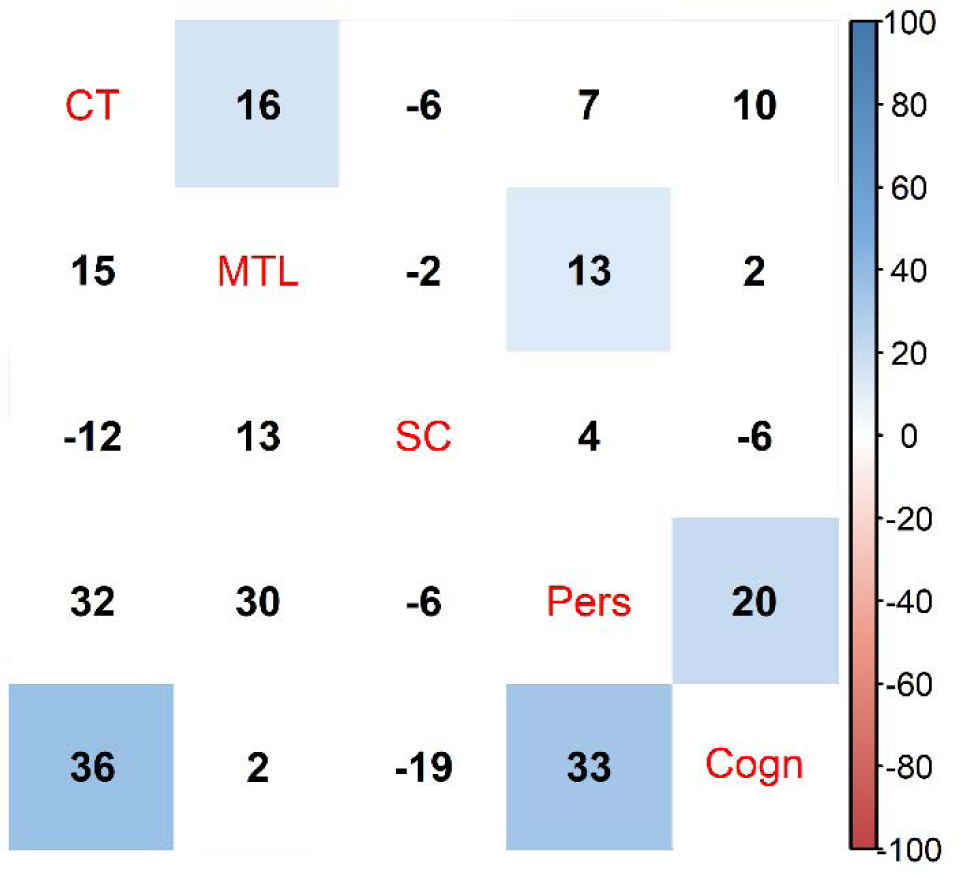
Phenotypic (upper triangle) and genetic (lower triangle) correlations between PPS-s used for heritability analysis. Phenotypic correlations account for family structure. FDR-corrected significant correlations are highlighted with color. Correlations are multiplied by 100 for clarity. Cogn=PPS of cognitive tests; corr=correlation; CT=PPS of cortical thickness; MTL=PPS of medial temporal lobe volume; Pers=PPS of personality tests; SC=PPS of subcortical structure volumes

**Fig. S12.**
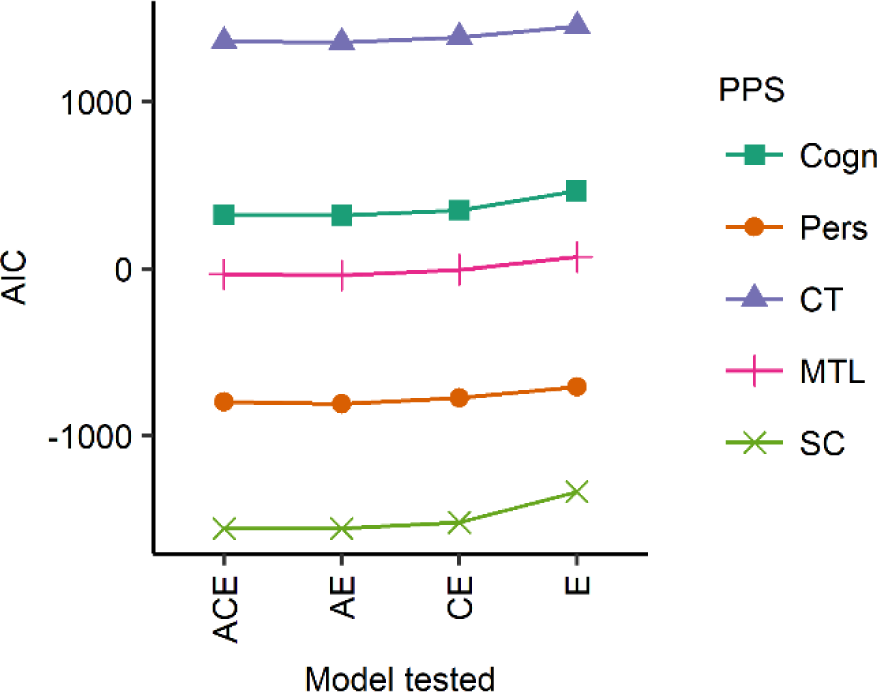
Akaike Information Criteria (AIC) for BMI-PPS (poly-phenotype score) bivariate heritability decompositions. Cogn=PPS of cognitive tests; corr=correlation; CT=PPS of cortical thickness; MTL=PPS of medial temporal lobe volume; Pers=PPS of personality tests; SC=PPS of subcortical structure volumes.

**Table S1.** Descriptive statistics of samples analyzed.

| Variable | S900 | S1200n | S1200n<br>unrelated |
| --- | --- | --- | --- |
| N | 895 | 225 | 124 |
| Age (years) | $\bar{x}=28.83$<br>(SD=3.67) | $\bar{x}=28.85$<br>(SD=3.84) | $\bar{x}=29.31$<br>(SD=3.83) |
| BMI (kg/m<sup>2</sup>) | $\bar{x}=27.27$<br>(SD=5.77) | $\bar{x}=26.51$<br>(SD=5.21) | $\bar{x}=26.32$<br>(SD=5.18) |
| BMI groups |  |  |  |
| Normal weight (BMI 18-24.9) | 375 (41.9%) | 101 (44.9%) | 56 (45.2%) |
| Overweight (BMI 25-29.9) | 285 (31.8%) | 74 (32.9%) | 45 (36.3%) |
| Obese (BMI 30+) | 235 (26.3%) | 50 (22.2%) | 23 (18.5%) |
| Drug test positive |  |  |  |
| No | 777 (86.8%) | 195 (86.7%) | 105 (84.7%) |
| Yes | 118 (13.2%) | 30 (13.3%) | 19 (15.3%) |
| Education (years) | $\bar{x}=14.85$<br>(SD=1.82) | $\bar{x}=15.06$<br>(SD=1.72) | $\bar{x}=14.83$<br>(SD=1.8) |
| Ethnicity: |  |  |  |
| Hispanic/Latino | 819 (91.5%) | 198 (88%) | 114 (91.9%) |
| Not Hispanic/Latino/unknown | 76 (8.5%) | 27 (12%) | 10 (8.1%) |
| Families | 384 | 151 | 66 |
| 1 sibling | 37 (10.4%) | 19 (20%) | 19 (28.8%) |
| 2 siblings | 107 (30.1%) | 49 (51.6%) | 36 (54.5%) |
| 3 siblings | 163 (45.9%) | 20 (21.1%) | 11 (16.7%) |
| 4 siblings | 43 (12.1%) | 6 (6.3%) | 0 (0%) |
| 5 siblings | 5 (1.4%) | 1 (1.1%) | 0 (0%) |
| Gender |  |  |  |
| Male | 413 (46.1%) | 120 (53.3%) | 61 (49.2%) |
| Female no birth control | 143 (16%) | 24 (10.7%) | 16 (12.9%) |
| Female with birth control | 339 (37.9%) | 81 (36%) | 47 (37.9%) |
| Handedness | $\bar{x}=65.07$<br>(SD=45.13) | $\bar{x}=68.93$<br>(SD=41.03) | $\bar{x}=70.73$<br>(SD=36.97) |
| Income |  |  |  |
| <\$10,000 | 65 (7.3%) | 16 (7.1%) | 9 (7.3%) |
| 10K-19,999 | 79 (8.8%) | 12 (5.3%) | 9 (7.3%) |
| 20K-29,999 | 116 (13%) | 24 (10.7%) | 15 (12.1%) |
| 30K-39,999 | 104 (11.6%) | 30 (13.3%) | 17 (13.7%) |
| 40K-49,999 | 98 (10.9%) | 23 (10.2%) | 13 (10.5%) |
| 50K-74,999 | 181 (20.2%) | 46 (20.4%) | 25 (20.2%) |
| 75K-99,999 | 119 (13.3%) | 28 (12.4%) | 14 (11.3%) |
| >=100,000 | 133 (14.9%) | 46 (20.4%) | 22 (17.7%) |
| Race |  |  |  |
| White | 664 (74.2%) | 176 (78.2%) | 95 (76.6%) |
| Other/unknown | 45 (5%) | 21 (9.3%) | 11 (8.9%) |
| Black or African Am. | 145 (16.2%) | 13 (5.8%) | 8 (6.5%) |
| Asian/Nat. Hawaiian/Other Pacific Is. | 41 (4.6%) | 15 (6.7%) | 10 (8.1%) |

BMI=body mass index; Is=islander; Nat=native

#### Additional Dataset S1 (separate file)

See first tab of file “SI_Dataset_1.xlsx” for table of contents.

